# MIMoSA: A Method for Inter-Modal Segmentation Analysis

**DOI:** 10.1101/150284

**Authors:** Alessandra M. Valcarcel, Kristin A. Linn, Simon N. Vandekar, Theodore D. Satterthwaite, Peter A. Calabresi, Dzung L. Pham, Russell T. Shinohara

## Abstract

Magnetic resonance imaging (MRI) is crucial for *in vivo* detection and characterization of white matter lesions (WML) in multiple sclerosis. While these lesions have been studied for over two decades using MRI technology, automated segmentation remains challenging. Although the majority of statistical techniques for the automated segmentation of WML are based on a single imaging modality, recent advances have used multimodal techniques for identifying WML. Complementary imaging modalities emphasize different tissue properties, which can help identify and characterize interrelated features of lesions. However, prior work has ignored relationships between imaging modalities, which may be informative in this clinical context. To harness the coherent changes in these measurements, we utilized inter-modal coupling regression (IMCo) to estimate the covariance structure across modalities. We then used a local logistic regression, MIMoSA, which leverages new covariance features from IMCo regression as well as the mean structure of each imaging modality in order to model the probability that any voxel is part of a lesion. Finally, we introduced a novel thresholding algorithm to fully automate the estimation of the probability maps to generate fully automated segmentations masks for 94 subjects. To evaluate the performance of the automated segmentations generated using MIMoSA we compared results with gold standard manual segmentations. We demonstrate the superiority of MIMoSA to other automated segmentation techniques by comparing it to the OASIS algorithm as well as LesionTOADS. MIMoSA resulted in statistically significant improvement in lesion segmentation.

## 1. Introduction

### 1.1 Multiple Sclerosis and MRI

Multiple sclerosis (MS) is an autoimmune disease of the central nervous system (CNS) that is characterized by pathologic changes in the brain and spinal cord. These pathologic changes include axonal injury and gliosis as well as demyelination, which is most prominent in focal white matter lesions (WML), although is also present in grey matter structures. It is well established that the accumulation of these WMLs is associated with disability and cognitive decline [1]. The in vivo assessment of lesion volume is primarily based on magnetic resonance imaging (MRI), as demyelination and other pathological changes cause tissue to have different water content compared to normal-appearing white matter [2]. The number and volume of lesions are essential metrics for monitoring disease progression in clinical settings, and are also used for evaluating the efficacy of disease-modifying therapies in clinical trials and in clinical practice [3].

The use of multiple MRI sequences can add significant value in identifying abnormalities in the brain. In MS, the most common MRI modalities acquired include; T2-weighted Fluid-Attenuated Inversion Recovery (FLAIR), T2-weighted (T2), Proton Density-weighted (PD), and T1-weighted (T1) images. WML appear as hyperintensities on the FLAIR, T2, and PD images while WML appear as isointense or hypointensities on the T1. The differing contrasts allow the viewer to detect different features of WML or normal-appearing white matter in order to delineate WML. For example, the FLAIR, unlike T2 and PD, easily distinguishes WML from cerebrospinal fluid (CSF) and thus is useful when evaluating lesions near CSF [4]. To gather as much information as possible about the demyelination occurring in the brain, it is now common to utilize the complementary information provided by different imaging sequences.

### 1.2 Lesion Segmentation (Manual, semi-automated, and automated methods)

Segmentation of WM lesions involves extracting locations in an image that contain white matter abnormalities, thus simplifying the image representation. Currently, manual segmentation is the gold standard approach in WML identification. Radiologists or other imaging scientists visually assess scans and manually delineate lesions on each slice in order to report total number and volume of WML. Not only is this costly and time-consuming, but it is prone to large inter- and intra- observer variability due to the challenge of incorporating 3D information from several MRI modalities [5] [6]. However, these WML metrics are vital in clinical trials where lesion number and volume are important outcomes for assessing disease-related changes and treatment effects [7]. In these clinical trials, a consistent method for quick and accurate delineation of WML is necessary. Though manual lesion segmentation is flawed, it has retained its primacy due to artifacts and errors that occur with automated and semi-automated methods.

Semi-automated methods cut some cost and time associated with manual lesion segmentation, but still require an imaging scientist to manually tune and verify segmentations. In a semi-automated procedure, an imaging scientist provides some input about their impression of the image, runs an algorithm, and obtains an initial estimate of lesion segmentation either as probability maps or a binary segmentation. The imaging scientist often then goes through and manually corrects the segmentation result. As semi-automated methods add a systematic component to the segmentation procedure they are less prone to inter- and intraobserver variability [8]. Nonetheless, similar to manual lesion segmentation, semi-automated methods are more costly and less timely than fully automated methods.

Automated methods eliminate the need for manual input, thus cutting cost and reducing implementation time even further. Automated methods additionally introduce stability and consistency into lesion segmentation as they eliminate human bias and error. Though many automated approaches and methods exist, no currently available algorithm is able to outperform manual lesion segmentation in terms sensitivity and specificity across subjects and scanning centers [5][8-9]. As a result, no particular automated segmentation algorithm is accepted as the gold standard in practice. Thus, accurate automated detection and delineation of WMLs remains a challenging unmet need in the field.

Most automated WML segmentation methods consist of two components: feature extraction and a classification algorithm. Classification algorithms range from sophisticated machine learning methods to simpler algorithms such as voxelwise logistic regression, linear discriminant analysis, and quadratic discriminant analysis [9]. Feature selection can also vary from simple raw intensities to complex functions of images. In the past, studies have compared classification methods, and shown that simple methods often yield performance equivalent to more sophisticated methods [10]. Such studies have pointed to the importance of biologically relevant feature selection [9]. This motivates the development of interpretable and discriminative features as key components for generalizable and accurate WML segmentation methods.

### 1.3 MIMoSA and IMCo Regression

We propose a fully automated segmentation algorithm that we refer to as a *Method for Inter-Modal Segmentation Analysis* (MIMoSA). As feature extraction is known to be pivotal for a segmentation algorithm’s accuracy and generalizability, we focus on the development of novel features. The majority of statistical techniques for the segmentation of WML are based on modeling intensity patterns for each imaging modality separately. However, recent advances in neuroimaging analysis have emphasized multimodal techniques in order to include covariance modeling across modalities [11] [12] [13]. This relationship, which we refer to as coupling or inter-modal coupling (IMCo), is known to differ across tissue types [14] [15]. However, it is unknown whether IMCo is disrupted in pathological conditions such as MS. We propose to leverage IMCo information as features for lesion segmentation in order to quantify the coherent changes as tissue damage and repair occur across imaging modalities. We introduce MIMoSA, a segmentation algorithm which utilizes inter modal covariance structures through harnessing our prior work regarding the coupling of different imaging parameters at a given anatomic location [16]. MIMoSA is a local-level logistic regression that accounts for mean structure as well as local covariance structure across imaging modalities. Additionally, we fully automate the model with a novel thresholding algorithm that detects the ideal threshold for probability maps in order to maximize similarity with gold standard manual segmentations. As described below, this approach is successful in detecting WML with increased accuracy.

## 2. Materials and Methods

In this section we introduce MIMoSA, a lesion detection method that harnesses IMCo analysis. Prior to fitting a local linear regression, we create a brain tissue mask excluding cerebrospinal fluid and extracerebral tissue. We intensity-normalize all MRI volumes, and identify candidate voxels that are hyperintense on FLAIR and thus contain the vast majority of white matter lesions. Additionally, we smooth volumes to capture the local spatial information while reducing artifact due to noise. We also utilize IMCo information in the model, which measures pathological changes across modalities. MIMoSA also includes a fully automated thresholding algorithm to create optimal binary lesion masks. MIMoSA captures the local covariance structure across scanning modalities through IMCo regressions at each voxel likely to contain lesional tissue. We then include IMCo regression intercept and slope estimates in the lesion classification procedure to obtain logistic regression coefficients, which we use to produce maps of the probability of lesion.

We evaluate the performance of MIMoSA on MRI volumes of the brain using a dataset collected at the Johns Hopkins Hospital consisting of 98 subjects with relapsing-remitting MS. We train the model and evaluate model performance using manually delineated segmentations. We train and test the proposed MIMoSA model and compare its performance to two competitively performing, previously published methods, OASIS [17] and LesionTOADS [16], by conducting bootstrapped cross-.

### 2.1 Study Population

We consider MRI studies from 98 subjects with MS. The median age of the MS subjects was 44 years (Q1, Q3) (33, 54), 72 are female, and the median EDSS was 3.5 (2, 6). Due to poor image quality, we excluded 4 subjects, which results in a total of 94 subjects.

### 2.2 Experimental methods

Local institutional review boards approved the imaging protocol and data analysis. 3D T1-MPRAGE (T1w) images (repetition time (TR) = 10 ms; echo time (TE) = 6 ms; flip angle (FA) a = 8°; inversion time (TI) = 835 ms, resolution = .828 mm × .828 mm × 1.1 mm), 2D T2-weighted FLAIR images (TR = 11,000 ms; TE = 68 ms; TI = 2800 ms; in-plane resolution = 0.83 mm × 0.83 mm; slice thickness = 2.2 mm), and T2-weighted (T2w) and PD (PDw) images (TR = 4200 ms; TE = 12/80 ms; resolution = 0.83 mm × 0.83 mm × 2.2 mm) were acquired on a 3T MRI scanner (Philips Medical Systems, Best, The Netherlands). Gold standard segmentations were acquired by an imaging technologist with more than 10 years of experience in delineating lesions and neuroanatomy.

### 2.3 Image Preprocessing

All images were preprocessed using the Medical Image Processing Analysis and Visualization (MIPAV) [18], TOADS-CRUISE [19], and Java Image Science Toolkit (JIST) software packages [20]. We first rigidly aligned the T1-weighted image of each subject into the Montreal Neurological Institute (MNI) template space at 1mm isotropic resolution. We used the normalized mutual information cost function for the co-registration and windowed sinc interpolation. We then registered the FLAIR, PD, and T2-weighted images of each subject to these aligned T1-weighted images. We also applied the N3 inhomogeneity correction algorithm [21] to all images and removed extracerebral voxels using SPECTRE [22].

### 2.4. Statistical modeling and spatial smoothing

We performed all statistical modeling in the R environment (version 3.1.0, R Foundation for Statistical Computing, Vienna, Austria) utilizing the packages oasis [23], ROCR [24], data.table [10], brainR [26], oro.nifti [27], and fslr [28]. We also used FSL for the three dimensional spatial smoothing of the volumes.

### 2.5. Brain tissue mask

The MIMoSA algorithm utilizes two masks for identifying tissue that may contain lesions: the brain tissue mask and the candidate mask [17]. We first identify voxels containing cerebral tissue but exclude cerebrospinal fluid (CSF). Because CSF appears hypointense on FLAIR, we exclude voxels with intensities below the 15th percentile after eliminating extracerebral voxels as detected by SPECTRE. We refer to this mask as the brain tissue mask. Since voxels within lesions appear as hyperintensities in the FLAIR volume, we restrict our classifier to exclude any voxels whose FLAIR intensities are not consistent with lesions: we select the 85^th^ percentile and above voxels in the brain tissue mask as candidate voxels. This step reduces computation time and restricts the modeling space, which we have found empirically reduces false positives.

### 2.6. Intensity normalization

As conventional magnetic resonance imaging volumes are acquired in arbitrary units, we use a statistical intensity normalization in order to model intensities across subjects. For this normalization we use a linear z-scoring method [29] [30] with the brain tissue mask, making the units of each modality easily interpreted as standard deviations of the variability across the brain.

### 2.7 Smoothed Volumes

To account for average signal intensities around each voxel, we use a sequence of Gaussian smoothers with varying kernel sizes to develop additional features that have been shown to aid in classification [17] [9]. These features have also been noted to mitigate missegmentation artifacts that are due to residual image inhomogeneities after N3 correction [17]. We use smoothed volumes with kernel parameters of 10 and 20 mm, which perform well empirically. Figure 1 shows an example of a smoothed volume for illustration.

**Figure 1:**
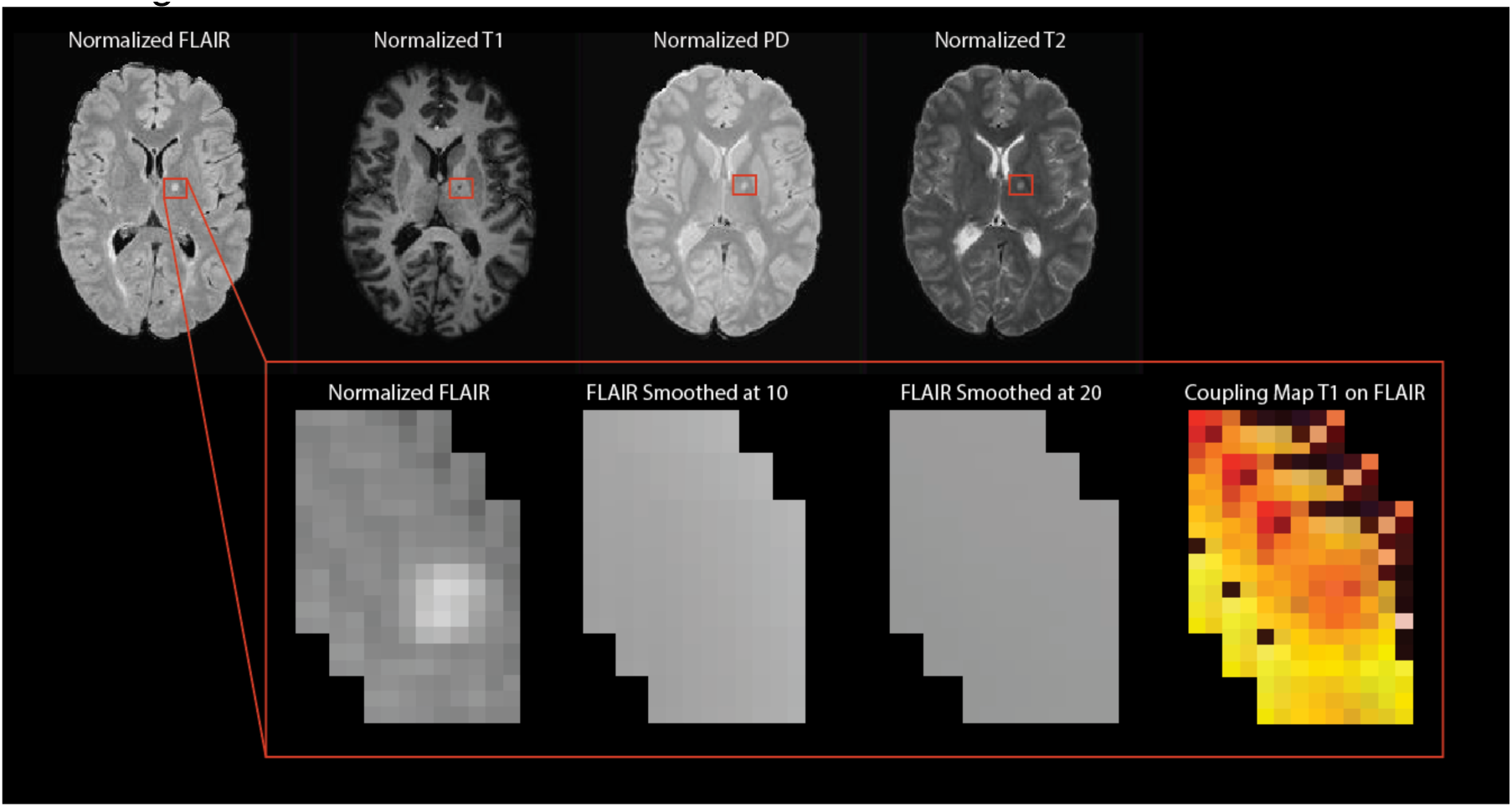
Features for MIMoSA including normalized images as well as an example of the FLAIR smoothed volumes and a coupling map for T1 on FLAIR IMCo regression.

### 2.8 Inter-Modal Coupling Regression

In order to help distinguish the lesional tissue from normal-appearing white matter, we utilize features we estimate from IMCo regression. These measures are intended to capture the local covariance structure across modalities as it varies across the brain. For example, as inflammation and demyelination occur in white matter lesions, not only do T1-weighted intensities decrease and FLAIR intensities increase; rather, voxels with more pathology tend to experience these changes concurrently to a greater extent. To quantify this, we perform a weighted local regression in a neighborhood around each voxel (see Figure 2), where the weight is proportional to a Gaussian kernel that is a function of the distance to the center voxel with fixed full-width half-max parameter (3mm) [11]. We record two coupling measures for each pair of imaging modalities at all voxels in the candidate mask: the locally estimated slope parameter as well as the intercept parameter estimate.

**Figure 2:**
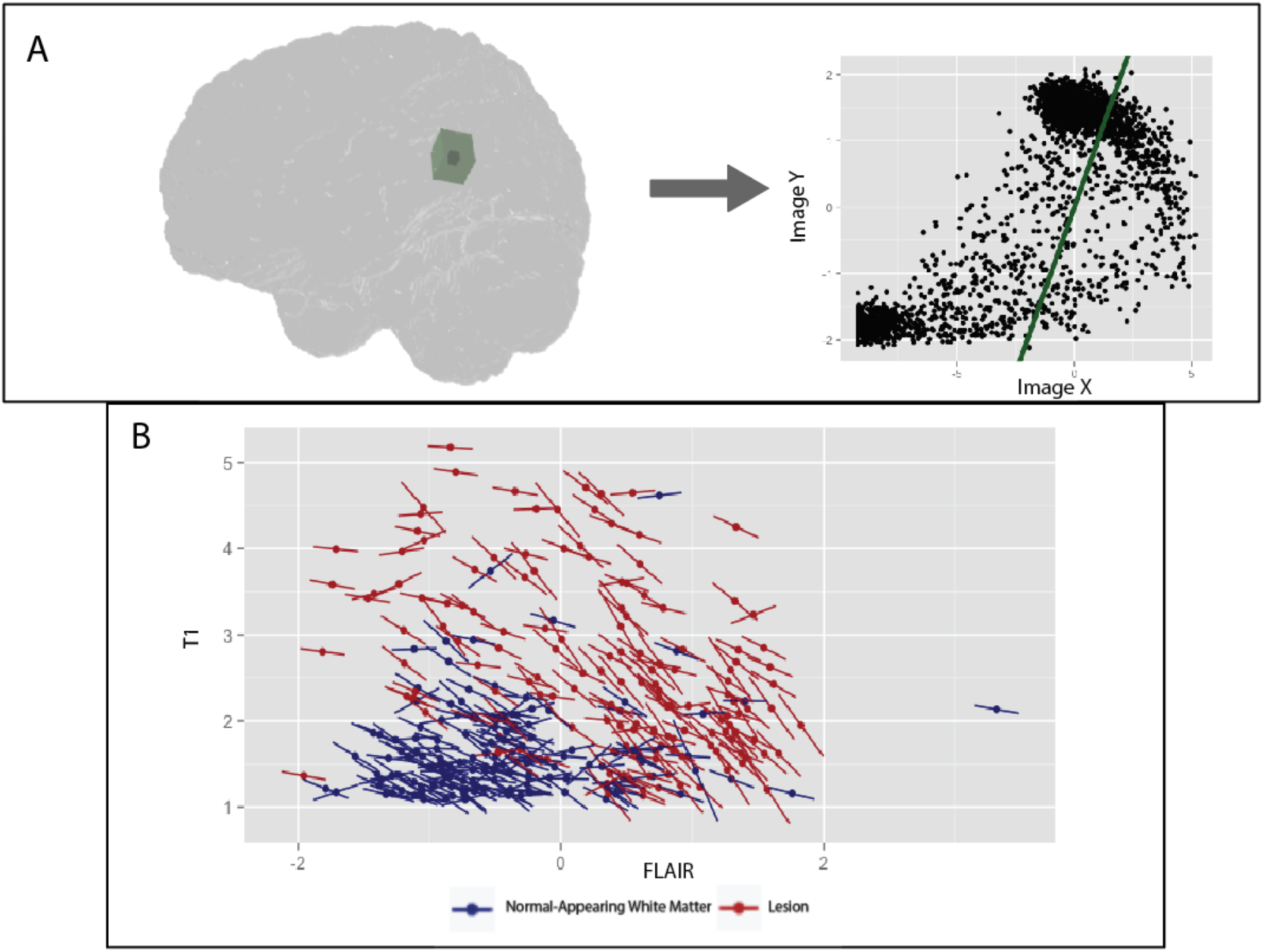
An example of IMCo regression for a single center voxel is shown in A. This is not to scale and solely for informational purposes. B shows a random sample of IMCo regression slopes and intercepts for full brain IMCo regression to emphasize the distinct pattern within lesion.

We implement IMCo regression on 12 pairs of inter-modal contrasts. That is, we exhaust all possible combinations (6) of the 4 scanning contrasts: T1 and PD, T1 and FLAIR, T1 and T2, PD and FLAIR, PD and T2, and FLAIR and T2. As IMCo is a regression, we must assign one modality to be the outcome and the other to be the predictor variable. Since there is no natural choice for predictor or outcome variables, for each pair we include both regressions which result in complementary information. For example, one combination of volumes is T1 and FLAIR, so we perform IMCo regression using T1 as the predictor variable and FLAIR as the outcome variable. We then repeat the IMCo regression with FLAIR as the predictor variable and T1 is the outcome variable. This leaves us with 12 unique pairs for estimating IMCo for which we obtain slope and intercept parameter estimates.

### 2.9 MIMoSA: A Method for Inter-Modal Segmentation Analysis

In this section we introduce the MIMoSA model. MIMoSA uses logistic regression to model the probability that a voxel contains lesional tissue. We choose logistic regression for two main reasons: first, it is relatively straightforward to interpret and implement. Second, a previous automated segmentation model showed promising results with a logistic regression compared with more advanced machine learning classifiers [9] but left significant room for improvement on the inclusion of intermodal features. We model the probability of lesion at the voxel level using FLAIR, PD, T2, and T1 intensities as well as the intensities from smoothed volumes of each modality (see Section 2.7). Using these features only captures the mean structure within modalities. For improved sensitivity and specificity to lesional tissue, we capture this covariance structure across scanning modalities using coupling measures for each pair of modalities, as described in the previous section, and we include these features in the model. Like all supervised lesion segmentation methods, we train MIMoSA on manually segmented images (see Figure 3).

**Figure 3:**
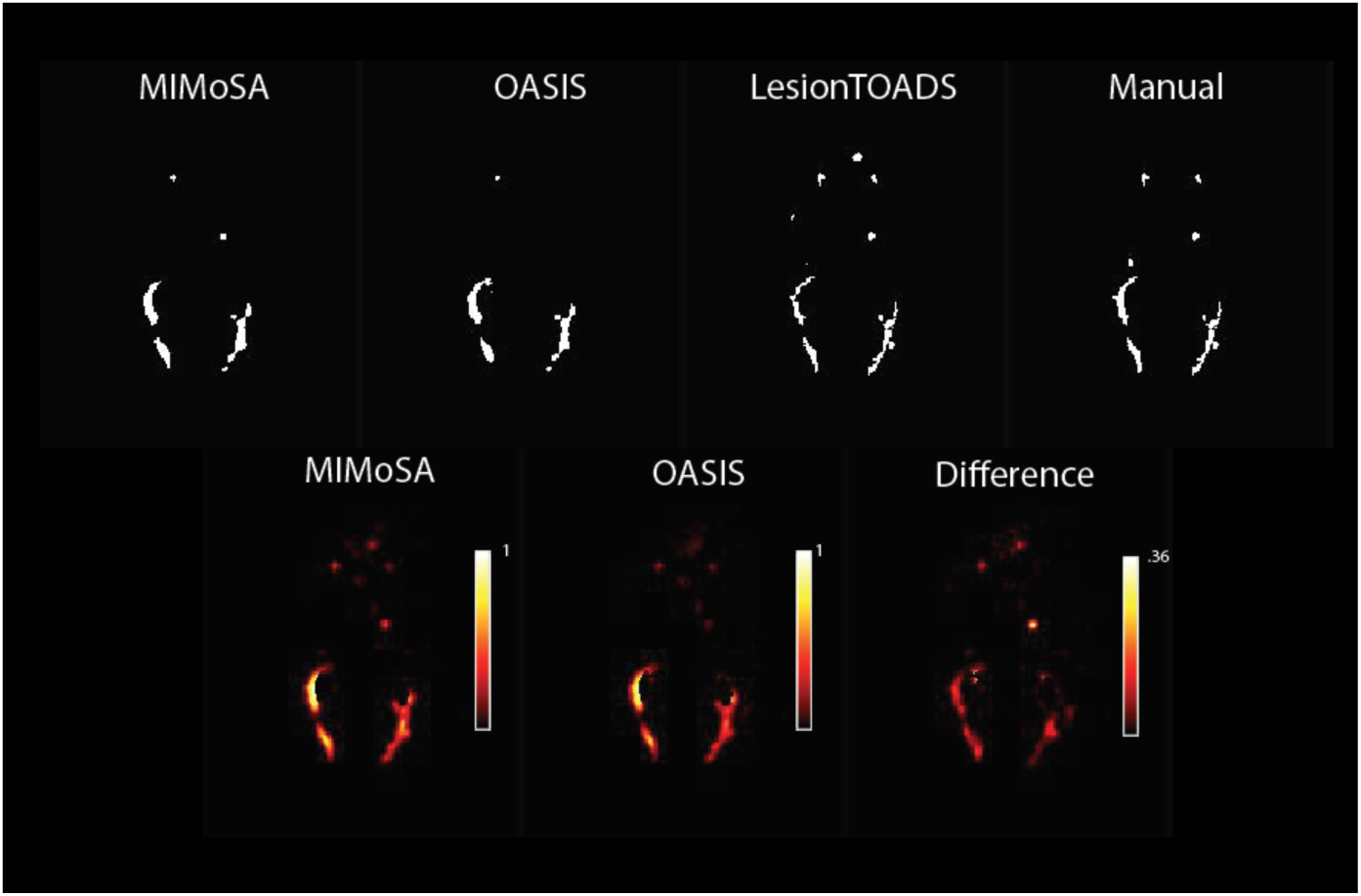
Probability maps for MIMoSA and OASIS as well as the difference (MIMoSA-OASIS) are shown in row 1. Using the thresholding algorithm lesion segmentations for respective models are also shown in row 2 along with LesionTOADS hard segmentations. LesionTOADS and gold standard manual segmentations are also shown.

The next step in the MIMoSA procedure is to fit a voxel-level logistic regression model over the candidate voxels. In the model below, *P*{*L_i_*(*v*) = 1} represents the probability that a voxel is part of a lesion where *L_i_*(*v*) is a random variable denoting voxel-level lesion presence. If there is a lesion in voxel *v* for subject *i*, then *L_i_*(*v*) =1, otherwise *L_i_*(*v*) = 0. We model the probability that a voxel *v* contains lesion incidence with the following logistic regression model:

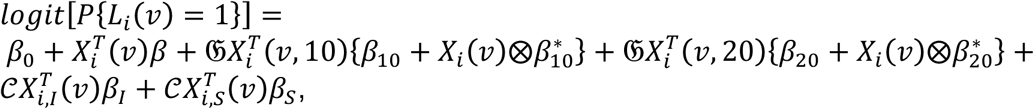

where we denote the normalized images *X_i_*(*v*) = [*T*_1,*i*_(*v*), *FLAIR_i_*(*v*), *T*_2,*i*_(*v*), *PD_i_*(*v*)]*^T^* and we use 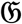 to denote the smoothing operator with parameter *δ* ∈ {10*mm*, 20*mm*}, which takes a weighted average within each neighborhood *N*(*v*, *δ*) around *v*. We express the smoothed images in vector form by 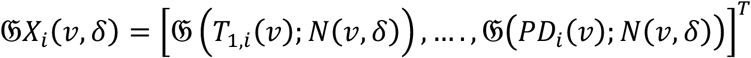, and we denote all combination of intercept and slope IMCo parameters respectively by 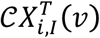 and 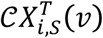. We use ⊗ to represent the Hadamard product. The interaction terms between the normalized volumes and the smoothed volumes, denoted by 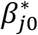, contribute to the model by capturing differences between voxel intensities and their local mean intensities. These aid in mitigating artifacts due to residual field inhomogeneity in some cases, and generally improve lesion detection performance.

After training, the result of our model is a collection of coefficients that can be used to determine the probability that each voxel is part of a lesion in a new set of images. MIMoSA obtains the estimated probability of each voxel being a part of a lesion by including the 12 imaging features (the four imaging modalities and the 2 smoothed volumes for each modality) and capturing the covariance across the 12 pairs of IMCo parameters. One can then apply a threshold to the probability maps to create binary lesion segmentation masks, which are typically preferred in clinical applications.

### 2.10 Probability map and binary segmentation

To determine where lesions are present, we estimate a probability map using the estimated regression coefficients for each voxel in the brain mask. We use a Gaussian smoother with sigma parameter of 1.25mm on this probability map to reduce noise. To create a binary segmentation, we use a population-level threshold on the smoothed probability map. Figure 3 shows a slice of a probability map and a binary segmentation after thresholding for a subject in a test set. In the next section, we describe an optimal threshold algorithm which allows the method to be fully automated by calculating a threshold yielding maximal overlap with the gold standard in the training set to create binary lesion masks.

### 2.11 Optimal Thresholding Algorithm

After the MIMoSA model is trained, it can be applied to generate probability maps, which are thresholded to create binary lesion segmentations. In comparable methods, this threshold is determined manually post hoc. To fully automate the thresholding process and select a threshold that maximizes similarity to gold standard manual segmentations, we introduce the optimal thresholding algorithm. The MIMoSA model is fit using the training set of data with gold standard manual segmentations. After the model is fit, we generate probability maps on these training set subjects. The optimal thresholding algorithm allows for the specification of a grid of thresholds. For each threshold we create binary lesion masks, which we compare to the gold standard manual lesion segmentations by calculating DSC. We select the threshold that produces the highest average DSC as the optimal threshold for application in the test set.

The optimal thresholding algorithm can be utilized in numerous ways. If investigators have a priori information about reasonable threshold values for their data they can simply create a finer threshold grid around the known value. If investigators are unsure about a reasonable threshold level a wider grid search can be utilized and then improved using finer grids. Results from the optimal threshold should include a variety of thresholds chosen from the interval as well as minimal thresholding at the boundary of the interval.

### 2.12 Bootstrap cross-validation

We conduct training and testing of both the OASIS and proposed MIMoSA methods using bootstrapped cross-validation. In order to fit the models and measure performance, we randomly allocate 42 subjects to the training set and 42 subjects to the test set. We then train both the OASIS and MIMoSA models using only subjects in the training set. After we fit the models, we apply the estimated coefficients to the test set in order to generate probability maps.

In addition to comparing MIMoSA with OASIS, we also compare against LesionTOADS, a segmentation algorithm based on fuzzy c-means that incorporates both topological constraints and a statistical atlas. Lesions are detected as outliers to the clustering functional. As LesionTOADS is an unsupervised learning method, we do not bootstrap the training and cross-validation. Instead, we simply apply the LesionTOADS algorithm with default parameters using the Java Image Science Toolkit [20] to obtain binary lesion segmentations for all 94 subjects directly.

In order to make the MIMoSA method fully automated, we propose a method to find the optimal threshold for binary lesion segmentation. We use this algorithm in conjunction with the MIMoSA method but we also apply it to OASIS for comparability. After each model is fit on a training set, we generate probability maps for the subjects in the training set. We then threshold these probability maps using values ranging from 20% and 35% in 1% increments to create hard segmentation masks. Using the set of predicted lesion masks for each model and each threshold, we calculate the Sørensen-Dice coefficient (DSC) at the subject level. After the DSC is calculated for each subject in the training set, we take the average across subjects for each threshold. We record the threshold with the highest average DSC score. We iterate this 100 times on each training set, and generate a frequency table of all optimal thresholds chosen by the algorithm. We apply the optimal threshold found in the training set for each iteration to the test set to provide binary lesion segmentation maps. We iterate this training and validation process to yield 100 bootstrap cross-validated sets of predicted probability maps and estimated binary segmentation masks.

It is not uncommon for MRI studies in MS to exclude collection of PD and/or T2. Due to this, we repeat the bootstrap cross validation procedure as if PD, T2, and both PD and T2 were not collected. This analysis does not only exclude the main effects but all possible interactions and IMCo measures. Additionally, we evaluate the method with the bootstrap cross validation when trained on only 20 subjects and tested on 74 subjects. These additional analyses evaluate the robustness of MIMoSA under different data collection schemes.

### 2.13 Calculation of summary statistics and confidence intervals

Using the cross-validated probability maps, we summarize performance results by subject-level partial area under the receiver-operator characteristic curve (pAUC, up to 1% false positive rate) [24] and DSC comparing the OASIS and proposed MIMoSA models. We use pAUC as it only considers regions of the ROC space which correspond to clinically relevant values of specificity [31]; that is, we do not consider model performance under clinically irrelevant high false positive rates. We then average performance measures across subjects and cross-validation folds. To compare performance statistically between MIMoSA and OASIS methods, we report confidence intervals for the difference between MIMoSA and OASIS (MIMoSA-OASIS) DSC and pAUC. To accomplish this, for each bootstrapped test dataset we first find and record the average difference in pAUC and DSC quantities for MIMoSA and OASIS. After averages are obtained, we find the values associated with upper and lower 0.025 quantiles to provide confidence intervals. We also record the frequency with which each threshold is chosen in the optimal threshold algorithm in order to compare the optimal thresholding for MIMoSA and OASIS methods. In order to compare the performance of MIMoSA with LesionTOADS we use the binary segmentations produced by the LesionTOADS algorithm to calculate the subject-level DSC and pAUC. To compare performance statistically between MIMoSA and LesionTOADS methods, we report confidence intervals for the difference between MIMoSA and LesionTOADS (MIMoSA-LesionTOADS) DSC and pAUC, averaged over each training set.

## 3. Results

### 3.1 Optimal Threshold Results

Table 1 shows the frequency of optimal thresholds chosen by the optimizing DSC algorithm for MIMoSA and OASIS models using threshold values of 20% to 35% by 1%. We find the optimal threshold with OASIS ranges from 24% to 28%. Within this range, we obtain a mode of 25%. MIMoSA utilizes a slightly wider spread of optimal thresholds ranging from 26% to 32%. Results for the replication of the bootstrap cross-validation when PD, T2, PD and T2 are excluded are similar and thus not provided here. For the cross validation with only 20 subjects in the training set, thresholds are also similar and thus are not provided here.

**Table 1:** Frequency of optimal thresholds for MIMoSA and OASIS across bootstrap iterations.

| Threshold | 24% | 25% | 26% | 27% | 28% | 29% | 30% | 31% | 32% |
| --- | --- | --- | --- | --- | --- | --- | --- | --- | --- |
| **MIMoSA** | 0 | 0 | 1 | 14 | 37 | 30 | 10 | 7 | 1 |
| **OASIS** | 13 | 25 | 30 | 16 | 6 | 0 | 0 | 0 | 0 |

In the OASIS algorithm, a recalibration of the population-level segmentation threshold was necessary and required manual adjustment. With the optimal threshold algorithm proposed here, this manual adjustment step is no longer required and allows fully automated segmentation of images from a new imaging center if training data are available.

### 3.2 Summary Statistics Results

Table 2 shows average DSC and pAUC across bootstrapped test samples and confidence intervals for the difference between MIMoSA and OASIS. MIMoSA outperforms OASIS and LesionTOADS in both average DSC and pAUC. The confidence interval for differences with OASIS do not contain 0, indicating that the observed improvement in both DSC and pAUC provided by MIMoSA are statistically significant. The confidence intervals for differences with LesionTOADS indicate a statistically significant difference in DSC but comparable pAUC.

**Table 2:** Average DSC and pAUC for MIMoSA, OASIS, and LesionTOADS as well as confidence intervals for their differences estimated using bootstrapped cross-validation.

|  | DSC | pAUC |
| --- | --- | --- |
| **MIMoSA** | 0.57 | 0.68 |
| **OASIS** | 0.54 | 0.63 |
| **LesionTOADS** | 0.50 | 0.65 |
| **MIMoSA - LesionTOADS (95% CI)** | (0.03,0.10) | (-0.01, 0.05) |
| **MIMoSA - OASIS (95% CI)** | (0.02,0.04) | (0.03,0.06) |

Table 3 shows average DSC and pAUC across bootstrapped test samples when PD and T2 images are excluded from analysis. Additionally, results are presented for bootstrapped samples where we train on 20 subjects and test on 74. When the model is trained using only 20 subjects we see negligible changes to average DSC, pAUC and associated confidence intervals. If a study is missing PD we see slight changes to DSC and pAUC while if T2 is excluded we see more moderate changes in these values. If both PD and T2 are excluded results are similar to when only T2 is missing. As we have shown previously that MIMoSA is statistically significantly better than OASIS and LesionTOADS table 3 does not show confidence intervals on the difference as each adaptation yields similar results. We additionally evaluate whether the variation in model yields results statistically significantly different from the partial models. Each interval contains 0 and thus there is not a loss in DSC or pAUC when these models are implemented.

**Table 3:** DSC and pAUC values for the MIMoSA models in replication of bootstrap cross-validation under exclusion of PD, T2, PD and T2, and training on 20 subjects.

|  | DSC | pAUC |
| --- | --- | --- |
| **No PD** | 0.56 | 0.66 |
| **No T2** | 0.55 | 0.65 |
| **No PD and T2** | 0.54 | 0.64 |
| **Training Set**<br>**20 Subjects** | 0.56 | 0.67 |

### 3.3 Qualitative Results

MIMoSA shows a significant improvement in performance over OASIS and LesionTOADS. In order to evaluate locations and patterns of differences between MIMoSA and OASIS, we compare probability maps. To compare with LesionTOADS, we compare binary segmentations from each respective algorithm. Figure 3 displays probability maps and binary segmentations from all models in axial slices while Figure 4 displays binary segmentation masks from all models in 3D allowing for global visualization of results. These results show that MIMoSA is able to identify lesions that OASIS and LesionTOADS were unable to detect. By inspection of the 3D visualizations in Figure 4, we note that MIMoSA performs better at juxtacortical lesion detection. Additionally, Figure 3 shows MIMoSA better separates lesions that are spatially close and which OASIS and LesionTOADS could not distinguish as distinct lesions. Furthermore, Figures 3 and 4 show OASIS and LesionTOADS tend to exhibit more false positive regions than MIMoSA, which is also reflected quantitatively in the ROC analysis.

**Figure 4:**
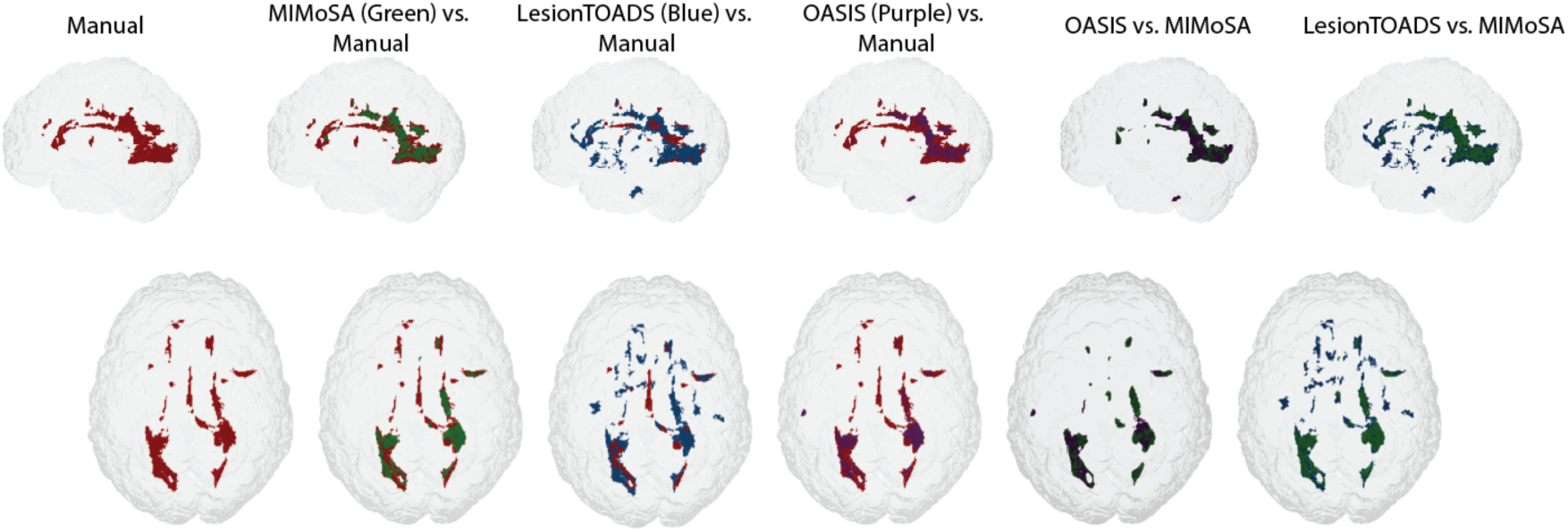
We compare 3D results in order to show globally segmentation performance of MIMoSA, OASIS, and LesionTOADS with gold standard manual segmentations.

## 4. Discussion

MIMoSA is a fully automated segmentation method that harnesses the changes in inter-modality covariance structure that occur in white matter pathology, and can be used to assist in WML detection or replace manual segmentation. The method is fully automated after training and does not require human input. MIMoSA avoids the introduction of human error cutting down on inter- and intravariability associated with manual and semi-automated WML segmentation. The model can be easily adapted and trained for cases when fewer imaging sequences are available. We show that MIMoSA yields superior segmentation results when compared to OASIS and LesionTOADS, which have been shown to be competitive in performance to the state of the art machine learning methods [28] [32].

Lesion segmentation methods, whether sophisticated machine learning classifiers or simpler methods such as intensity-based regression models, depend on the development and refinement of discriminative reliable imaging features. IMCo regression aims to detect biological changes reflecting processes occurring inside WML captured in the different scanning modalities. IMCo modeling facilitates new opportunities for feature extraction for the purpose of WML segmentation, but also promising new measures of pathological severity and repair processes [33]. These results are shown to be robust when fewer subjects are available to train, and if certain imaging modalities such as PD and T2 are not available. As such, IMCo regression features could be useful in not only for volumetric analyses but also hold promise for monitoring disease and quantifying effects of disease-modifying therapies.

For generalizations to data from different imaging centers or protocols, the recalibration of a threshold can be achieved automatically and optimally using the proposed cross-validation scheme. This novel algorithm estimates the best threshold for probability maps for producing segmentations that maximize similarity with gold standard manual segmentations. MIMoSA can thus easily be applied in a fully automated manner to new datasets when gold standard segmentations on training subjects are available.

Future work includes further validation of MIMoSA under variations in imaging protocols in order to show the replicability of IMCo measures and segmentation performance. Additional opportunities for performance improvement may also include the refinement of IMCo regressions for lesion segmentation by including complex modeling of nonlinear IMCo relationships, as well as the use of multiple neighborhood sizes in multi-scale IMCo analyses. An investigation can also be done on whether all pairwise combinations of modalities is necessary using model selection procedures. Furthermore, MIMoSA could be a useful tool in lesion segmentation in longitudinal studies but should be evaluated under different training schemes to ensure validity. Beyond binary segmentation maps, the method shows promise in providing information about WML with different intermodal information that might aid in the adjudication of causes of lesions, for example comparing vascular to demyelinating contributions. Moreover, IMCo regression coefficients can be useful features in longitudinal studies for modeling prediction of lesion behavior and progression.

## Acknowledgements

The authors would like to thank John Muschelli who provided valuable feedback concerning the software implementation of coupling. Additionally, we would like to thank Elizabeth Sweeney for providing useful feedback. The project described was supported in part by a pilot grant from the Center for Biomedical Computing and Analytics at the University of Pennsylvania as well as RO1NS085211, R21NS093349, R01EB017255, R01MH107703, R01NS082347 from the National Institutes of Health. Additionally, this project was supported in part by NMSS grant RG-1507-05243. The content is solely the responsibility of the authors and does not necessarily represent the official views of the funding agencies.

